# Pharmacoinformatics based elucidation and designing of potential inhibitors against *Plasmodium falciparum* to target importin α/β mediated nuclear importation

**DOI:** 10.1101/2020.12.29.424688

**Authors:** Arafat Rahman Oany, Tahmina Pervin, Mohammad Ali Moni

## Abstract

*Plasmodium falciparum*, the prime causative agent of malaria, is responsible for 4, 05,000 deaths per year and fatality rates are higher among the children aged below 5 years. The emerging distribution of the multi-drug resistant *P. falciparum* becomes a worldwide concern, so the identification of unique targets and novel inhibitors is a prime need now. In the present study, we have employed pharmacoinformatics approaches to analyze 265 lead-like compounds from PubChem databases for virtual screening. Thereafter, 15 lead-like compounds were docked within the active side pocket of importin alpha. Comparative ligand properties and absorption, distribution, metabolism, excretion, and toxicity (ADMET) profile were also assessed. Finally, a novel inhibitor was designed and assessed computationally for its efficacy. From the comparative analysis we have found that our screened compounds possess better results than the existing lead ivermectin; having the highest binding energy of −15.6 kcal/mol, whereas ivermectin has −12.4kcal/mol. The novel lead compound possessed more fascinating output without deviating any of the rules of Lipinski. It also possessed higher bioavailability and the drug-likeness score of 0.55 and 0.71, respectively compared to ivermectin. Furthermore, the binding study was confirmed by molecular dynamics simulation over 25 ns by evaluating the stability of the complex. Finally, all the screened compounds and the novel compound showed promising ADMET properties likewise. To end, we hope that our proposed screened compounds, as well as the novel compound, might give some advances to treat malaria efficiently in vitro and in vivo.

## 1.0 Introduction

Malaria, a disease caused by the infection through protozoan parasites of the genus *Plasmodium,* disseminates to humans by female anopheles mosquitoes and is responsible for about 228 million medical cases in recent years. It is also accountable for nearly half a million deaths, and the severity rates are higher among the children (Carter & Mendis 2002), (Snow *et al.* 2005). Among the five species of malaria, *P. falciparum* is the deadliest and responsible for 99.7% of malaria cases in the African region. That is considered as a hot-spot of malaria infection, having 213 million or 93% cases in that region while the total worldwide cases were 228 millions.(Organisation 2018).

The ever-rising distribution of the multi-drug resistant *P. falciparum* now becomes a global concern and is mediated by the resistance to conventional antimalarials, chloroquine and artemisinin(Ashley *et al.* 2014; Takala-Harrison *et al.* 2015; Coppée *et al.* 2020). As a consequence, there is a pressing necessity to develop novel antimalarial agents that are effective against drug-resistant malarial parasites and might be working on different or unique targets and also structurally distinct from existing antimalarial agents (Peters *et al.* 1990; Palmer *et al.* 1993)

The unambiguous contrasts in gene control components standing in the nucleus between *P falciparum* and the human could prompt new potential drug targets for antimalarial drug development(Wittayacom *et al.* 2010).

Nuclear transport also shows a foremost role in disease conditions such as oncogenesis and viral illness (Hogarth *et al.* 2006; Yoneda-Kato *et al.* 2008; Ao *et al.* 2010). The periphery between nuclear and cytoplasmic compartments is infiltrated by NPC that permit swapping of macromolecules between the nucleus and cytoplasm. On the other hand, nuclear import facilitates higher molecular mass proteins through a selective and energy-dependent manner with the aid of NLS and NES. Importin family proteins identify these targeting sequences that expedite the nuclear import of their cargo proteins mediated by the formation of importin α/β heterodimer(Fontes *et al.* 2000).

Precise inhibitors of nuclear import symbolize valuable tools for scrutinizing nuclear vehicle technology in forthcoming research. Leptomycin B, an antifungal antibiotic, has already been tested and is a well-characterized compound for inhibiting nuclear transport. Ratjadones and some small-molecule inhibitors have also been suggested for its efficacy to inhibit nuclear importation(Kudo *et al.* 1999; Köster *et al.* 2003).

Ivermectin has also been suggested as a promising candidate for inhibiting nuclear import at a precisely lower concentration and thereby killing the pathogens. It is also very selective to importin α/β mediated import and having no effects on other importin family proteins(Chaccour *et al.* 2013; Panchal *et al.* 2014).

In the present study, we have tried to design a novel inhibitor of this unique target by using pharmacoinformatics, bioinformatics techniques for drug designing, approaches to combat against deadly malaria.

## 2.0 Materials and Methods

The overall proposed molecular mechanism for drug designing is shown in Figure 1.

### 2.1 Sequence retrieval

The NCBI database was initially explored to find out *P. falciparum* importin α and β proteins (Wheeler *et al.* 2007). Protein importin α (gi|29501524|) and β (gi|29501526|) were selected for the study. The sequences were then stored as a FASTA format for further analysis.

### 2.2 Homology modelling, structural refinement and model validation

Homology models of the targets were done by MODELLER v9 (Sali *et al.* 1995),and top-scoring template was selected for the model construction. The templates used for the modeling were 4TNM_A and 3W3W_A for importin α and β, respectively. The probability and the identity scores for importin α were 100% and 49% and for β-100% and 26%, respectively. The structural refinement of the models were done by GalaxyWEB server (Ko *et al.* 2012). Then the predicted models were employed for the validation through the PROCHECK (Laskowski *et al.* 1996)tool of the SWISS-MODEL Workspace (Arnold *et al.* 2006). Additionally, the ProSA-web server (Wiederstein & Sippl 2007) was also utilized for assessing the model quality through Z-score measurement.

### 2.3 Electrostatic potential analysis and energy minimization

The electrostatic potential of the predicted models was built by APBS (Unni *et al.* 2011)server through transforming.pdb to .pqr file and the visualization was done by PyMOL (version 1.3) molecular graphics system (DeLano 2002). For further securing the validation of the predicted models, the energy minimizations were also done. The GROMOS 96 (Scott *et al.* 1999) force field was used for the minimization of the structure and validated by the “Ramachandran Plot” statistic by using the SWISS-MODEL Workspace (Arnold *et al.* 2006).

### 2.4 Active site analysis and virtual screening

The active sites of the final proposed models were constructed by the CASTp server (version 3.0). This server builds a model through the pocket algorithm of the α shape theory (*Dundas et al. 2006*). The screening was initially performed based on existing literature to get an insight into the potential inhibitors, and then the PubChem database (Kim *et al.* 2016) was screened for similar hits. All the ligands selected for the docking studies were converted into the. pdbqt format using the Open Bable toolbox (O’Boyle *et al.* 2011). Autodock Vina (version 1.1.2) (Trott & Olson 2010) was used for the screening. The grid generated for molecular docking at the center was X: 35.09, Y: 18.24, and Z: − 12.06. The dimensions used for the docking analysis were X: 51.00, Y: 40.00. and Z: 39.00.

### 2.5 Ligand properties and ADMET analysis

For the cross-validation of the ligands as an efficient therapeutic candidate, ligands structural and behavioral properties were analyzed.. Wide ranges of properties were evaluated including Lipinski’s Rule of Five (Lipinski 2004), drug-likeness, molecular weight, and bioavailability. These parameters were assessed through various online servers and tools including, Molinspiration (http://www.molinspiration.com), Drug-likeness tool (Oprea 2000), and OSIRIS Property Explorer (Sander 2001). The absorption, distribution, metabolism, excretion, and toxicity (ADMET) were also assessed for the selected compounds. The ADME SARfari (Davies *et al.* 2015), admet- SAR (Cheng *et al.* 2012), Swiss database (Wirth *et al.* 2013), and QikProp (3.5) (Schrödinger 2012) were used for these assessments.

### 2.6 Novel lead design

The novel lead was designed based on the screening outcome with the assistance of PubChem Sketcher V2.4 (Ihlenfeldt *et al.* 2009) through the previously mentioned method (Oany *et al.* 2014a; Oany *et al.* 2020b, a). Since the screened compounds and the ivermectin itself violated few properties of Lipinski’s rules for drug designing, we have further analyzed to design a novel compound to eradicate the limitations. To design the novel leads, we initially selected the PubChem compounds C_31_H_44_O_7_ and C_32_H_48_O_8_ as base templates. We have chosen the best candidate retaining the core ring structure of ivermectin following extensive *in silico* trial and error, including drug-likeness, bioavailability, and molecular docking. The designed structure was then fetched as Canonic SMILES format. Then the three-dimensional structure was generated by the Open Babel toolbox (O’Boyle *et al.* 2011). The Ligand properties, molecular docking, and ADMET analysis of the novel compounds were also assessed as described earlier.

### 2.7 Molecular dynamics Simulation

The molecular dynamic approach is widely used to assess atoms’ behavior, structural stability and to study the conformational changes on the atomic level. Here we applied the molecular dynamics simulation for the designed compound by nanoscale molecular dynamics (NAMD) (V2.1) software (Phillips *et al.* 2005), and the temperature was set to 303 K. Configuration files for MD simulations were generated by CHARMM-GUI website (http://www.charmm-gui.org/) (Jo *et al.* 2008). Ligand parameterization was performed using CHARMM General Force Field (CGenFF) web-based tool (https://cgenff.umaryland.edu/). The energy was minimized for 10000 steps, and production for 25 ns timescale under the NPT ensemble.

## 3.0 Results

### 3.1 Homology modelling, structural refinement and model validation

Homology models of the importin α and β were constructed through MODELLER and are illustrated in **Fig. 2a and 2b**. These models are visualized by the PyMOL (version 1.3) molecular graphics system. The GalaxyWEB server refined the models with the best poses and finally, the validations of these models were tested by constructing the “Ramachandran Plot” and its statistics (**Fig 2a**, **2b**,and **Table S1**). Furthermore, the ProSA Z score was also implemented for validation and shown in **Fig 2a, 2b**,subsequently. The combined view of importin α and β is shown in **Figure 2c.**

### 3.2 Electrostatic potentiality and energy minimization

The electrostatic potentiality of the modeled importin α and β proteins were analyzed for the identification of the energy distribution of the protein and showed in **Fig. 3a and 3b**. For better model construction, we applied the energy minimization process onto the modeled importin α and β proteins through the GROMOS 96 force field. The force-field energies of the modeled proteins before and after minimization were 6114.219 and −32996.316 kJ/mol for importin α, whereas −6404.763 and −16,370.318 kJ/mol for importin β, respectively.

### 3.3 Active site exploration and virtual screening

CASTp server (version 3.0) predicted different active sites of our desired proteins with different volume scores and we picked the best large volume as the final active site (shown only for importin α) (**Fig. 4a**). The molecular surface area of the active site was 19480.291 Å. Finally, both structures were combinedly assessed for the common active site (**Fig. 4b**). A total of 265 lead-like compounds from over thousands of compounds (**Table S2**) were initially screened from the PubChem database based on mother compound Ivermectin as potential inhibitors of importin α through applying the filters (Molecular Weight: < 700; Rotatable bond count: <6; Heavy atom count:< 50; H-bond donor count: <4; Xlogp: <4 and Polar area, Å²:< 140). Thereafter based on drug-likeness and Lipinski’s rules, we have finally selected **15** lead-like compounds for docking, including Ivermectin (**Fig. S1**). Docking stances with 0.0 RMSD were selected as the final binding energies and interacting amino acid residues are illustrated in **Fig. 5**. The docking integration analysis results are shown in **Table 1**.

### 3.4 Ligand properties and ADMET data analysis

Different properties of the predicted small molecules (ligand) including drug-likeness, Lipinski’s rules, aqueous solubility, and bioavailability were assessed for the identification of the best therapeutic candidates (**Table 2**). All the ADMET properties were also assessed for the selected ligands and cross-check with different servers and tools for the validity of the results. These properties are described in **Table 3**.

### 3.5 Novel drug design

The designed novel drug from PubChem Sketcher V2.4 is shown in **Fig 6a**, having the molecular formula of C_25_H_37_O_8_. The ligand properties of the novel drug are shown in **Table 2.** Interaction analysis from the molecular docking is illustrated in **Fig 6b** and within the active site pocket is Fig **6c**. The ADMET profile is described in **Table 3**. The amino acid residues involved in docking for the novel compound are also shown in **Table 1**.

### 3.6 Molecular dynamics Simulation

To understand the stability, flexibility, structural behavior, and binding mechanism of the best protein-ligand complex along with the apo form of protein, 25 ns of molecular dynamics simulations were conducted. Here, we considered our designed novel compound, with the importin α protein. The atomic RMSDs of the protein-ligand complex along with the importin α protein were calculated and are plotted in a time-dependent manner in Figure 7a, where the importin α protein and protein-ligand complex are indicated with the yellow and blue color, respectively.

According to Figure 7a, these two indexes, i.e. the apo form of protein and the protein-ligand complex, regularly fluctuate around a constant value and become stabilized considerably from 17 ns to end. To understand the flexible region of the protein and the movement of each residue, the RMSF value of the amino acid sequence of the respected protein complex was explored along with the single protein and is depicted in Figure 7b. The protein-ligand complex showed slightly higher fluctuations from 270-330 residues compared with the single protein.

## 4.0 Discussion

Nuclear importation is very crucial for the survival of the eukaryotes and is mediated by protein, importins (Wälde & Kehlenbach 2010). *P. falciparum* possesses only a pair of importins-importin α and β within its genome (Mohmmed *et al.* 2005). Lack of auto-inhibition of importin α is a unique feature of apicomplexan parasites including *P. falciparum.* It might have a role in the faster growth of the parasites (Dey *et al.* 2018). The present study focuses on this unique target for designing a novel small molecule inhibitor.

The formation of importin heterodimer α/β is required for the nuclear importation and is mediated by NLS through its binding with importin α. Then importin β binds with importin α for facilitating the docking to the nuclear envelope and abetting translocation of the cargo protein into the nucleus (Görlich *et al.* 1999). Homology modeling showed both the importins with satisfactory values for the most favored regions of the Ramachandran plot having 95.36% and 95.43% for importin α and β, respectively. The Z scores from the ProSA server for the importin α and β were −10.48 and −14.0, which also support the validity of the predicted models, as the values were within the plot and close to zero. The active site predicted from the CASTp server (version 3.0) of importin α (**Fig 4a**) and the docked α and β showed (**Fig 4b**) us the striking phenomenon that the importin β docked within the active site of importin α. These would lead us to design small molecule inhibitors targeting this active site to inhibit the formation of heterodimer α/β. Application of energy minimization through GROMOS 96 force field steps also prepares the model ready to dock organization on its large active site pocket (Oany *et al.* 2018).

Ivermectin, the known inhibitor of nuclear import (Wagstaff *et al.* 2011; Wagstaff *et al.* 2012) had also been studied on *P. falciparum* (Panchal *et al.* 2014). As ivermectin possesses some characteristics that hamper its full potentiality as a lead compound, hence we are looking for better novel therapeutics to inhibit importin α as well as the formation of heterodimer α/β.

As per Lipinski’s rule of five for potential lead candidates (Lipinski 2004), ivermectin has failed to possess three major properties. The molecular weight should not be higher than 500 M, no more than 10 HBA, and partition coefficient (xlogP) no more than 5.0, and ivermectin had the values for the three parameters 875.1 M, 14, and 5.53, respectively. Besides, the complexity score of ivermectin was also much higher, which was 1680 (Kim *et al.* 2016).

From the 265 lead-like screened compounds, based on the mother compound ivermectin, **13** compounds (**Table 2**) met the properties of Lipinski’s rule of five though slightly deviating the molecular weight, ranging from 528 to 584 M. The molecular docking simulation of the selected compounds had shown very promising results compared to ivermectin (−14.6 kcal/mol) and ranged from −13.0to −15.6 kcal/mol. The bioavailability scores of the selected compounds were 0.55 and ivermectin as well as doramectin had only the value of 0.17. The drug-likeness score of ivermectin was inspiring in comparison to other screened compounds. The ADMET analysis of all the screened compounds as well as ivermectin and doramectin revealed that they possessed similar profiles (**Table 3**). Ivermectin and doramectin both had lower water solubility and intestinal absorptions. The PubChem ID 6476306, 10209711, and 11226920 possess P-glycoprotein I inhibitor activity along with ivermectin and doramectin.

We found that there were few limitations for all the screened compounds as well as ivermectin itself to become a potential lead candidate; for these reasons, we tried to design a novel lead compound to overcome all these limitations (Oany *et al.* 2014b; Hasan *et al.* 2016; *Hossain et al. 2016;* Oany *et al.* 2017). Initially, we designed about 15 compounds (not shown here) and from them, the novel compound with a molecular formula of C_25_H_37_O_8_ was selected for final analysis. The molecular weight, partition coefficient (xlogP), and the HBA value were 465.25, 0.98, and 8, respectively. The binding energy for this compound was −12.4 kcal/mol, and it is impressive compared to its molecular weight. The Arg 325 residue was found to be very important from the interaction analysis with ivermectin through creating hydrogen bond and the designed lead compound also formed a hydrogen bonding with the Arg 325 residue. Additionally, both the ivermectin and designed lead also formed steric interaction with Asn 1025 amino acid residue. These could be very much promising for the inhibition of dimer formation. The drug-likeness score for the novel compound (0.71) was similar to ivermectin and had a promising bioavailability score of 0.55. The binding groove for the novel compound was also similar to ivermectin and within the active site pocket. The aqueous solubility (log S) value for the novel compound was −3.35. That means the substance is soluble in water, whereas the value for poorly-soluble ivermectin was −9.70. Finally, the MD simulation study of the targeted protein and the novel designed compound further reinforced our prediction by validating the complex interaction stability presented by the RMSD value (Figure 7a). The RMSF values of the single protein and the complex have also been observed, and few higher fluctuations were detected in the looped regions of the protein (Figure 7b).

In conclusion, from the comparative analysis, we are quite optimistic that the screened compounds along with the novel compound might put roles as therapeutic compounds over ivermectin.

## 5.0 Conclusion

Nucleo-cytoplasmic shuttling is very crucial for the survival and growth of malaria parasites. The ever-rising strains of malaria have recently become a global concern, and researchers are currently looking for more effective targets for designing better functional drugs. Hence, importins could become a very promising target for further potent drug designing, and more extensive experimental works are required to conduct in this sector. Nowadays, pharmacoinformatics approaches are quite promising for designing novel therapeutics. These processes also help us to design therapeutics in a cost and time convenient way. Therefore, it is concluded that the designed novel compound might be exploited further for its efficacy in vitro and in vivo. Nevertheless, our findings led researchers to work extensively on this target for developing novel therapeutics.

## 6.0 Author contributions

Arafat Rahman Oany conceived, designed, and guided the study; drafted the manuscript; and analyzed the data. Tahmina Pervin carried out the analysis, drafted the manuscript. Mohammad Ali Moni participated in coordination, performed critical revision, and helped in drafting the manuscript. All authors read and approved the final manuscript.

## 7.0 Acknowledgements

We thank Dr. K. M. Kaderi Kibria, Associate Professor, Department of Biotechnology and Genetic Engineering, Faculty of Life Science, Mawlana Bhashani Science and Technology University, Tangail, Bangladesh for helping us to design the work with his expertise.

## 10.0 Reference

